# Genetic characterization of the Zika virus epidemic in the US Virgin Islands

**DOI:** 10.1101/113100

**Authors:** Allison Black, Barney Potter, Gytis Dudas, Leora Feldstein, Nathan D Grubaugh, Kristian G Andersen, Brett R Ellis, Esther M Ellis, Trevor Bedford

## Abstract

Here we release draft genome sequences of Zika virus (ZIKV) that were sequenced from PCR-positive diagnostic specimens collected by the United States Virgin Islands Department of Health (USVI DoH) as part of their ongoing response to the ZIKV outbreak. We will use these sequences to conduct a genomic epidemiological study of ZIKV transmission in the USVI. We are releasing these genomes in the hope that they are useful for those individuals involved in the public health response to ZIKV and to other groups working to understand Zika virus transmission and evolution.

## Purpose

This work represents a pre-publication sharing of pathogen genomic data of public health significance. We believe that open sharing of sequence data and early analyses is necessary to inform public health response in a timely fashion [1]. We encourage other investigators to use these data in their own analyses to better contextualize their work. All we ask is that you please let us know if you plan to use these data in a publication. We intend for this announcement to serve as a ‘marker paper’ specifying analyses we have planned for these data, namely to investigate Zika transmission in the USVI. For greater detail please see the ‘Genome Announcement’ below. We plan to replace this ‘announcement’ with a preprint and eventually with a publication. The citation will be continually updated throughout this process.

## Genome Announcement

For pathogens causing primarily asymptomatic infections, such as ZIKV, traditional epidemiological methods may yield biased estimates of incidence and outbreak growth due to incomplete case counts. Additionally, contact tracing to infer transmission chains and geographic patterns of spread may also break down when the majority of cases are unrecorded. Genomic epidemiological methods make use of a fundamental principle that the evolution of pathogens occurs on the same timescale as hosts transmit pathogens. Viral replication results in the accumulation of mutations, evidence that an infection has occurred even if asymptomatic or unreported. Thus using sequence data, in combination with temporal and demographic data, we can date when an outbreak started, describe the spatial spread of disease, infer how quickly an outbreak is growing, or determine transmission chains at the individual or the population level.

In collaboration with the US Virgin Islands Department of Health (USVI DoH) we are sequencing Zika virus (ZIKV) full genomes from PCR-positive diagnostic specimens collected during the ongoing DoH response to the ZIKV outbreak. We extracted viral RNA from inactivated serum or urine samples using a QIAamp Viral RNA mini kit (Qiagen). Viral RNA was reverse-transcribed, amplified, and sequenced according to the protocol in Quick et al [2]. Genomes were sequenced on either the MinION or Illumina MiSeq, as indicated, with a subset of samples sequenced on both platforms to ensure the validity of the sequences across platforms.

Our study seeks to describe when ZIKV was introduced to the USVI, how many separate introductions of ZIKV occurred, the degree to which different introductions contributed to local transmission, and describe patterns of ZIKV spread between the different US Virgin Islands. We will also use joint inference of genomic and epidemiological data to estimate the proportion of cases that are asymptomatic, and use inferred case counts to more accurately estimate *R*_0_ over the outbreak.

## Preliminary results

To date we have sequenced 11 Zika viruses from the USVI. In Figure 1 we show where they fall in the broader spectrum of viral diversity. It is clear that many of the infections are quite similar, while other infections appear to represent shorter transmission chains following introduction.

**Figure 1.**
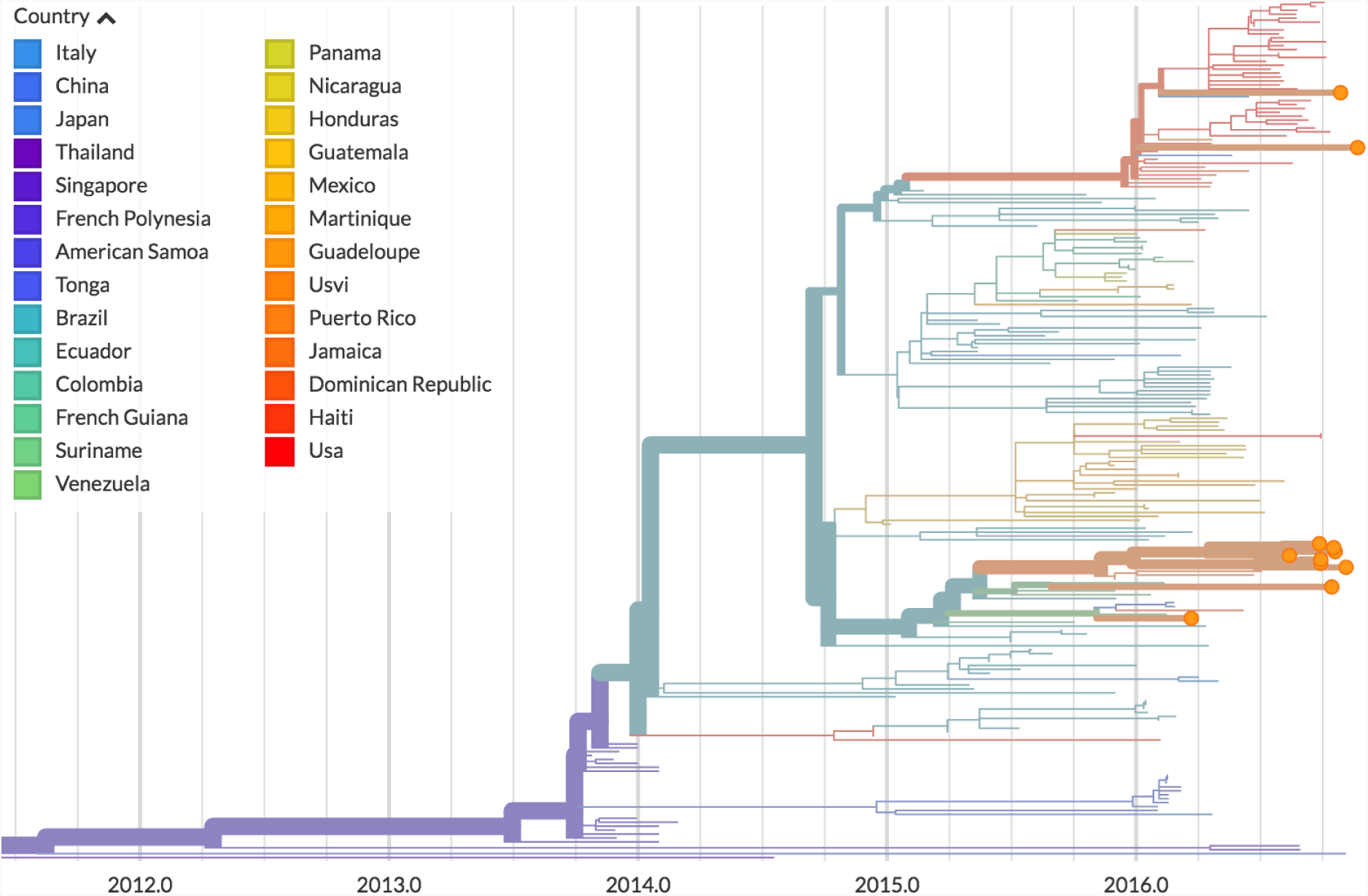
Preliminary phylogeny showing 11 Zika genomes from the USVI. This tree is taken from nextstrain.org/zika and shows a time calibrated phylogeny of publicly available Zika genomes.

## Data

All genome sequence data and metadata can be found on GitHub at github.com/blab/zika-usvi.

